# Contrasting biotic and abiotic drivers of Glomeromycotina and Mucoromycotina mycorrhizal associations in a durum wheat field

**DOI:** 10.64898/2026.02.18.706634

**Authors:** E. Taschen, E. Guillot, C. Plassard, E. Kerbiriou, D. Dezette, A. Taudière, A. Personne, A. Robin, D. Redecker, P. Hinsinger

## Abstract

Mucoromycotina fine root endophyte (M-FRE), although commonly present in cultivated crops, represent a largely overlooked symbiosis, and their diversity and ecological functions under natural field conditions remain poorly understood. The co-occurrence of M-FRE and Glomeromycotina arbuscular mycorrhizal fungi (G-AMF) was assessed in field-grown durum wheat (*Triticum turgidum* subsp. *durum* (Desf.) Husn.), testing the effects of combined water and nitrogen limitation on root colonization and fungal community diversity in roots, rhizosphere, and extra-radical hyphae.

The M-FRE colonization was reduced under combined water and nitrogen stress but was unaffected by wheat genotype. In contrast, G-AMF colonization varied among genotypes and was insensitive to this combined stress. While G-AMF colonization correlated with root traits, M-FRE abundance was rather determined by soil properties and the applied stress. Remarkably, under stress, M-FRE but not G-AMF colonization correlated with nitrogen and phosphorus uptake in plant shoots. Partial 18S metabarcoding detected 74 G-AMF taxa and 12 M-FRE taxa, some shared across compartments, revealing active growth of M-FRE extra-radical hyphae. Stress had contrasting effects on diversity: G-AMF alpha diversity remained stable, whereas M-FRE diversity declined, with stress driving distinct community structures for both groups.

These results indicate that M-FRE and G-AMF are shaped by divergent drivers, highlighting functional differences between these morphologically similar symbioses.

## Introduction

Mycorrhizal fungi are among the most widespread and functionally important root symbionts, forming mutualistic associations with the majority of terrestrial plants (Smith & Read, 2010). These symbioses enhance plant nutrient acquisition, particularly phosphorus (P) and nitrogen (N), improve water uptake, and confer increased tolerance to both abiotic and biotic stresses (Smith & Read, 2010). In agricultural systems, arbuscular mycorrhizal fungi (AMF) can contribute to sustainable production by enhancing nutrient use efficiency and maintaining soil health (Rillig *et al.,* 2019; Thirkell *et al.,* 2017). In Mediterranean regions, simultaneous water and nitrogen limitations are major constraints for crop growth and yield, and crops will increasingly face combined water and nutrient shortages in the coming decades (MedECC 2020). Mycorrhizal fungi can modulate plant responses to such combined stresses by improving nutrient acquisition and water relations, yet their own activity and effectiveness are also strongly influenced by environmental limitations, including N and water availability (Tang *et al.,* 2022).

Most research on AMF has focused on the Glomeromycotina subphylum (G-AMF), which are well known for their intra-radical arbuscules and extraradical hyphae that facilitate nutrient exchange (Smith & Smith, 2011). More recently, the Mucoromycotina fine root endophytes (M-FRE) have emerged as an ecologically significant but understudied lineage (Orchard *et al.,* 2017ab; Field *et al.,* 2018), following recent phylogenetic reclassification within the Mucoromycotina (Orchard *et al.,* 2017a) subphylum, notably the establishment of the genus *Planticonsortium* (Walker et al., 2018) for the fine root endophyte *Glomus tenue*, now called *Planticonsortium tenue*. Arbuscules formed by M-FRE appear functionally similar to those of G-AMF as described for former *Glomus tenue* (Gianinazzi-Pearson et al., 1981), and revealed by cryo-scanning electron microscopy and X-ray microanalysis (Albernoz *et al.,* 2022). These structures contribute to plant acquisition of phosphorus (P) and nitrogen (N), as demonstrated in liverworts (Field *et al.,* 2019) and in a flowering plant species (*Trifolium repens*; Hoysted *et al.,* 2019). Although M-FRE share some functional traits with G-AMF, their distinct evolutionary histories raise questions about their physiology and ecological strategies (Prout *et al.,* 2019), particularly in agricultural systems, as M-FRE exhibit marked affinities for disturbed environments (Albernoz *et al.,* 2021, 2022). The recent analysis of the assembled metagenome of a putative M-FRE confirms higher saprotrophic abilities than Glomeromycotina fungi such as the capacity for degradation of microbial cell walls, and a complete cellulose degradation pathway (Cole *et al.,* 2025). This raises fundamental questions about their respective responses to environmental filters, their roles in plant nutrition and ecosystem functioning, and consequently their importance in agroecosystems.

Mycorrhizal colonization and community composition are strongly influenced by abiotic factors such as nutrient availability, soil moisture, and temperature, as well as by biotic factors including plant genotype and root traits (Smith & Read, 2010, Powell & Rillig, 2018). Nutrient availability, especially P, is known as a major driver: high P supply often reduces AMF colonization, but the response varies according to Glomeromycotina species (Wakelin *et al.,* 2012, Trinquier *et al*. 2026), and to sub-phyla (Glomeromycotina and Mucoromycotina; Jeffery *et al.,* 2018). Biotic factors such as host identity and genotype also modulate mycorrhizal interactions, influencing both colonization levels (Ganugi, *et al*. 2021) and G-AMF community structure of close species (*Medicago sp*.; Pivato *et al.,* 2007) and microbial communities among wheat varieties (Jacquiod et al,. 2022). These drivers may interact in complex ways, with potential consequences on the functioning of M-FRE and G-AMF symbioses.

Glomeromycotina fungi interact with plants within the roots and extend their extraradical hyphae into the surrounding soil, particularly beyond the nutrient depletion zone created by roots. These hyphae can explore up to 10 cm away from root surface, depending on the G-AMF species, considerably improving the access to nutrients with low mobility, such as P (Thonar *et al.,* 2011). Some degree of compartment-specific strategies has been documented, with intra-radical structures primarily mediating nutrient exchange and potentially contributing to plant protection, while extraradical hyphae mainly facilitate nutrient acquisition (Powell *et al.,* 2009) and soil aggregation (Smith & Smith, 2011; Maherali & Klironomos, 2012). Only a few studies have addressed phylogenetic trait conservatism and the evolution of functional trade-offs in arbuscular mycorrhizal fungi (Hart & Reader, 2002; Powell *et al.,* 2009). The hyphae of M-FRE also grow out of roots, as shown on the only cultivated isolate existing to date (Howard *et al.,* 2024), and produce small extra-radical hyphae and very small spores able to act as propagules for plant inoculation (Thippayarugs *et al.,* 1999). But, to date, it is unknown whether all M-FRE species exhibit similar or distinct patterns of compartment-specific colonization compared with G-AMF. Understanding the diversity and localization of M-FRE structures is essential to decipher the functional roles of these fungi in agroecosystems, and their functional diversity and complementarity with G-AMF.

This study aims to advance our ecological understanding of these two types of mycorrhizal symbioses by comparing how G-AMF and M-FRE respond to abiotic and biotic factors under field conditions. Focusing on durum wheat (*Triticum turgidum* subsp. *durum* (Desf.) Husn.), we investigated their responses to environmental stress and host genotype, and measured plant N and P uptake to explore potential correlations with both fungal types. We further examined community composition and diversity across three compartments (roots, extra-radical hyphae, and rhizospheric soil) to reveal compartment-specific patterns. By combining microscopic quantification of colonization with high-throughput metabarcoding, this study provides a direct comparison of G-AMF and M-FRE ecological responses, offering new insights into their regulation and potential functional complementarity in agroecosystems.

## Material & Methods

The studied field was located in Southern France (°38’48.0"N, 3°52’59.1"E) at the experimental site of the GEVES North of Montpellier. This study site is characterized by deep clay–limestone loamy soils, with mean concentrations of 13 mg Olsen P kg⁻¹ soil, 0.92 g total N kg⁻¹ soil, and 95 g total C kg⁻¹ soil with 8.1 g organic C kg⁻¹ soil. Additional soil physicochemical properties are provided in **Table S1.** Fifteen durum wheat genotypes (**Table S2**) were cultivated, including varieties from Italy, the USA, and France, including the two most widely grown French varieties, Anvergur and RGT Voilur, which account for 55.7% and 9.4% of the national durum wheat surface area, respectively (Arvalis, 2025), as well as a genotype from the experimental Evolutionary Pre-breeding population described by David *et al*. (2014). The field was divided in two halves (0.28 ha each) to apply two treatments: a control with optimal nutrient and water supply (151 kg N ha^-1^, two irrigations of 20 mm each) and a combined water and N stress treatment (110 kg N ha^-1^, a single irrigation of 25 mm). Hereafter, these treatments are referred to as Control and Stress. The field had previously been cultivated with soybean (*Glycine max*). Wheat genotypes were cultivated on plots of 1.5 x 2.5 m, separated by a buffer of a common variety (RGT Voilur). Each genotype was repeated on two plots per treatments (**Suppl. Fig. 1**). Each plot consisted of eight rows sown with a planting density of 325 seeds. m^-2^. Seeds were sown at the end of October 2020. In January 2021, one herbicide treatment was applied at growth stage 2 nods (GS 32; BALI at 0.15 dm^3^.ha^-1^). Final sampling was conducted by mid-April 2021. The phenological stage of plants was close to heading stage (variable according to varieties). On each plot, two samplings were done on the opposite side of the plot leading to a total of four samples for each genotype for each treatment. Plants were uprooted using a spade-fork over a soil volume of approximately 12 dm³ (0.2 m × 0.2 m × 0.3 m depth). The rhizosphere, i.e. operationally defined at the soil adhering to roots, was sampled by gently brushing the roots, then sieved at 2 mm, and stored at 4°C for nutrient analyses and -20 °C for DNA extractions. Roots were washed and separated from the aboveground part by cutting at the base of the stem, and separated into three subsamples, one for DNA extraction (after being blotted on a paper towel and stored at -20°C), another one for root morphology measurements (kept moist and then stored at -20°C), and the last one for mycorrhizal colonization analyses (stored in 70% alcohol at 4°C). Hyphal mesh bags were installed by mid-January 2021, two per plot, as described in the sampling design. Each bag was made of nylon mesh (37-µm pore size, preventing root ingrowth; dimensions 6.5×9.5 cm), sewn with nylon thread to avoid decomposition in the soil. The bags were filled with 130 g of river sand (particle size 0.5–2 mm). Holes for installing the hyphal mesh bags were made using a soil corer (8 cm diameter, 15 cm depth; **Suppl. Fig. 1**). The hyphal bags were sampled at the same time as the final sampling, by mid-April 2021.

### Root morphology

Roots were scanned in water-filed, transparent trays at 800 dpi and images were analyzed with the WinRHIZO software (2021; Regent Instruments, Quebec city, Canada). The root subsamples were then oven-dried at 60 °C for 2 days and weighed to determine the dry mass. The specific root length (SRL) was determined by dividing the total root length of the subsample by its dry mass, the root tissue density (RTD) by dividing the dry mass of the subsample by the volume obtained with WinRHIZO. The average diameter was determined automatically and used to calculate the fine root proportion (FRP) for those roots with a diameter < 0.4 mm.

### Mycorrhizal colonization in roots

Roots were stained following the ink and vinegar method (Vierheilig *et al.,* 1998). Briefly, roots were incubated overnight (14 h) at room temperature in 10% KOH (a milder approach than hot KOH for fragile roots), rinsed in water and then in water acidified with 8% vinegar. Samples were subsequently incubated in an 8% ink solution (Schaeffer® black permanent ink diluted in 8% acetic acid) for 15 min at 70 °C in a water bath, and rinsed three times with vinegar-acidified water. Stained roots were then bleached in glycerol-acetic acid for 30 min and stored in a vinegar–water solution at 4 °C until microscopic observation.

Thirty root fragments of 1 cm were observed at 100x to 400x for AMF colonization assessment according to an adapted method of Trouvelot *et al*. (1986) evaluating global mycorrhizal colonization and specific colonization and arbuscules intensity of G-AMF and M-FRE structures (Taschen *et al*., in prep). To determine the type of fungus, careful observation at 200x magnification was carried out, sometimes requiring 400x magnification to distinguish the closely intertwined hyphae, and a thorough examination of the arbuscules, observing the diameter and appearance of the hyphae to identify the fungus (Tippayarugs *et al.,* 1999; Orchard *et al.,* 2017ab; Prout *et al.,* 2024). M-FRE hyphae were characterized by their thin diameter (≈1 µm, compared to ≈10 µm for G-AMF), their darker staining, and their irregular, chaotic trajectories within root tissues, and the formation of arbuscules. In contrast, G-AMF hyphae typically follow long, straight intercellular pathways and intracellular colonization with arbuscules (ranging from Arum to Paris morphologies; Dickson *et al.,* 2004), spreading from cell to cell, with vesicles or intra-radical spores positioned at hyphal termini. Intraradical M-FRE hyphae were further distinguished by fine-angled branch points (fan-shaped ‘carrefours’) as well as by the formation of intercalary vesicles.

#### Soil nutrient analyses

At least 15 g of rhizosphere were collected by gently brushing roots and sieving the soil through a 2-mm mesh. Mineral N was extracted from 10 g of fresh soil with 50 cm^3^ of 1 M KCl (1 h shaking) and assayed for NH₄⁺ and NO₃⁻ using a flow injection analyzer (Continuous Flow Analyzer, Skalar). Available P (Olsen P) was measured on air-dried soil following Olsen method, extracting 2.5 g of soil with 50 cm^3^ of 0.5 M NaHCO₃ (pH 8.5) for 30 min. Phosphate in the extracts was quantified colorimetrically using the malachite green–oxalate method.

#### Plant N and P content

Plant numbers were recorded, and shoots were harvested, weighed, dried at 55 °C, and finely ground. Shoot total N content was determined by dry combustion (Thermo Flash 2000; ThermoFisherScientific). Shoot P content was measured after nitric acid mineralization (HNO₃ 65%, Milestone ETHOS EASY microwave) using a colorimetric assay: phosphate was quantified with the yellow vanadomolybdic method. Total N and P uptake per plant was calculated by multiplying shoot concentration by shoot dry mass.

### AMF diversity

#### DNA extraction

Root DNA was extracted using hot MATAB lysis with RNase treatment followed by magnetic-bead purification on a KingFisher™ Flex (program PEX-EXT-002q). Full protocol is provided in Supplementary Materials (**Suppl. doc 1**). Soil DNA was extracted using NucleoSpin® Soil and following the manufacturer’s indications.

#### Metabarcoding

For metabarcoding, the first PCR step was done as triplicates using the AMF specific primer pair AMV4.5NF/AMDGR (350 bp; Sato *et al*. 2005), firstly used for metabarcoding by Suzuki *et al.,* 2020 and which have been shown to partially amplify Mucoromycotina by Orchard *et al*. (2017a). The reaction conditions were as follows: initial denaturation at 95 °C for 3 min; 35 cycles of 95 °C for 30 s, 55 °C for 30 s, and 72 °C for 30 s, and a final elongation step at 72 °C for 5 min. All PCR amplifications were checked on agarose gels to verify successful amplification before pooling. The pooled PCR1 amplicons were purified by magnetic beads (Clean PCR, Proteigene, France). The second PCR was performed using a Nextera® XT Index Kit (Illumina, San Diego, USA) following the manufacturer’s instructions. After purification with magnetic beads, these final PCR products were multiplexed and sequenced on a MiSeq Illumina sequencer using MiSeq Reagent Kit v3 (600-cycle; Illumina).

#### Sequence processing and taxonomic assignment

All analyses were performed in R (code available as a target plan in <u>github</u>) using packages targets (Landau 2021), dada2 (Callahan *et al*. 2016), phyloseq (McMurdie and Holmes 2013) and MiscMetabar (Taudière 2023) packages. In short, primers were removed using *cutadapt* (https://cutadapt.readthedocs.io/en/stable/) and low-quality sequences were removed using the dada2::filterAndTrim function (with default parameter values). Amplicon sequence variants (ASVs) were inferred, forward and reverse reads were then merged, and sequences shorter than 200 bp as well as chimeras were discarded.

Each ASV was taxonomically assigned using the dada2::assignTaxonomy function, with the EUKARYOME SSU database as reference (v. 1.9.3 Tedersoo *et al*. 2024). The ASVs were post-clustered into OTUs (Operational Taxonomic Units) as recommended by Tedersoo *et al*. (2022) using function MiscMetabar::asv2otu; by then, they were called OTU. Further refinement was performed using the MUMU software (<u>v</u>.1.0.2 https://github.com/frederic-mahe/mumu) which re-clusters redundant or erroneous sequences.

From this taxonomic assignment, we retained OTUs belonging to the Glomeromycota. For Mucoromycotina, we filtered for the families *Endogonaceae* and *Densosporaceae*. However, since the precise taxonomic identity of arbuscule-forming Mucoromycotina remains uncertain, we applied a second approach: Mucoromycotina OUT were retained if they showed ≥95 % identity using BLAST algorithm against M-FRE reference sequences from previous studies, hence Mucoromycotina OTU suspected to be M-FRE. These references represented root endophytes isolated from a range of plant hosts such as bryophytes (liverworts, mosses), lycophytes, ferns, and early-diverging fungal lineages (*Endogone* spp.) and sampled across Europe, Africa, Asia, and Oceania. Sequences originated from NCBI from the following: Bidartondo *et al*. 2011 (JF accessions), Desirò *et al*. 2013 (KC accessions), Orchard *et al*. 2017a (KX accessions), and Rimington *et al*. 2015 and 2019 (respectively KJ and MH accessions).

#### Phylogeny

For phylogenetic reconstruction, we included the above-listed sequences and used Glomeromycotina as the ingroup, with *Coprinus comatus* (AY665772.1) and *Schizosaccharomyces pombe* (X54866.1) as outgroups. Additional reference sequences included *Rhizophagus irregularis* D-E1 (KU136417.1), uncultured *Acaulospora* OByBM VTX00023 clone GAMF5821 (LN621100.1), *Claroideoglomus sp*. 3 LB-2012 clone 3 from black chernozem soil (JX301671.1), and *Acaulospora laevis* WUM46/W3107 clone pWD95-1-4 (Y17633.2). All sequences were aligned with **MAFFT v7.525** (Katoh *et al.,* 2002) using the options --auto --adjustdirection --reorder. The aligned dataset was analyzed with IQ-TREE v3.0.1 (Wong *et al.,* 2025) where the best-fit model is automatically selected by ModelFinder (best model: TIM2+F+I+R3) and we computed ultrafast bootstrap approximation using option -B 1000.

### Statistical analyses

All analyses were performed in R (v. 4.3.0).

#### Colonization rates

Root colonization abundance of G-AMF and M-FRE was arcsine square-root transformed prior to analysis. Linear mixed-effects models were fitted using the lmer function (package lme4), with fungal type, treatment, and their interaction included as fixed effects. Plot identity (defined as genotype within spatial block) was included as a random effect to account for the non-independence of multiple samples collected within the same plot. Model significance was assessed using likelihood ratio tests, and post hoc comparisons were performed using estimated marginal means (emmeans package). Model assumptions were verified by inspection of residuals.

To identify which plant root traits and soil variables influenced AMF colonization rate (for G-AMF and M-FRE separately), we used a model selection approach with the glmulti package in R. Mycorrhization (M%) by G-AMF and M-FRE expressed as percentages, were arcsine-transformed prior to analysis. Root colonization by each fungal type was modeled as a function of root traits (specific root length (SRL), fine root proportion (FRP), root tissue density (RTD), root biomass), rhizosphere characteristics (available P (Olsen P), KCl-extractable ammonium and nitrate), colonization of the other fungal type and the stress treatment. Linear models (lm) were fitted, and model selection was based on corrected Akaike Information Criterion (AICc), considering only main effects (level = 1) and retaining the top 50 models. Best-supported models were those within ΔAICc ≤ 2 of the lowest AICc, and model-averaged adjusted R² values were calculated to estimate overall explanatory power. Residuals of the top model were checked for normality using Shapiro–Wilk tests and QQ plots. Parameter estimates, unconditional confidence intervals, and model importance were extracted for interpretation. Correlations between root colonization and plant N and P uptake were tested using Pearson correlation. The same approach was applied to examine correlations between root colonization and sequence abundances of G-AMF and M-FRE.

#### Metabarcoding data

Sequencing depth varied strongly among compartments, averaging 27,314 reads for roots (range: 2,007–48,497), 10,081 for hyphae (59–34,435), and 14,043 for soil (807–49,522). Mucoromycotina–Endogonaceae accounted for 14.7%, 4.0%, and 11.1% of reads in roots, hyphae, and soil, respectively.

All sequences were initially considered to obtain a global, descriptive overview of fungal diversity across sample types, i.e. compartments (roots, hyphae mesh bags and soil), with respect to relative abundance at the family level. Due to the low replication per genotype (n = 4), genotype effects could not be reliably assessed and were therefore excluded from fungal community analyses. The analyses focused on spatial compartmentalization of taxa and their responses to stress treatments. Quantitative diversity analyses require normalization of sequencing depth and were performed on filtered and rarefied datasets. Diversity analyses were performed separately for G-AMF and M-FRE. For G-AMF, samples with fewer than 1,000 reads were excluded, resulting in 119 root, 117 soil, and 23 hyphae samples. For M-FRE, only root and soil samples were retained due to low amplification in hyphae, yielding 99 root and 56 soil samples. Within each dataset, samples were rarefied to the minimum sequencing depth using phyloseq::rarefy_even_depth() prior to diversity analyses.

Alpha diversity (Hill numbers) was calculated with the renyi function from the *vegan* package, and Welch’s t-tests (*stats* package) were used to assess the effect of Stress treatment. Hill–Shannon and Hill–Simpson indices were interpreted as sensitive to differences in the relative abundance of common species (Roswell *et al.,* 2021).

Beta diversity was assessed separately for G-AMF and M-FRE using PCoA on Bray–Curtis distances (phyloseq::ordinate,), with PERMANOVA (MiscMetabar::adonis_pq) testing differences between sample types and stress treatments.

## Results

### Impact of wheat genotype, nutrients and root morphology on root colonization

Durum wheat root were colonized by both G-AMF and M-AMF fungi, forming typical and distinctive structures (**Fig. 1**). Arbuscules of both fungi occurred at close vicinity, in neighboring cortical root cells. Global (undistinguishing G-AMF and M-FRE) wheat root colonization was 16.8 %, and was not significantly impacted by stress treatment (p= 0.16) but varied according to wheat genotypes (p= 0.001). G-AMF root colonization did not respond to stress treatment (mean root colonization was 10.9 % in the stress treatment and 9.7 % in the control treatment; **Table 1**; **Fig. 2**) but M-FRE colonization was significantly affected (mean colonization: stress 4.5 %, control 8.0 %; **Table 1**; **Fig. 2**). Under control conditions, root colonization by both G-AMF and M-FRE was not significantly correlated with plant P or N uptake. Under stress, M-FRE colonization was positively correlated with plant nutrient uptake (for P, r = 0.40, p= < 0.001; for N, r = 0.33, p = 0.01), whereas G-AMF colonization remained uncorrelated with N or P (**Suppl. Fig. 2**).

**Figure 1.**
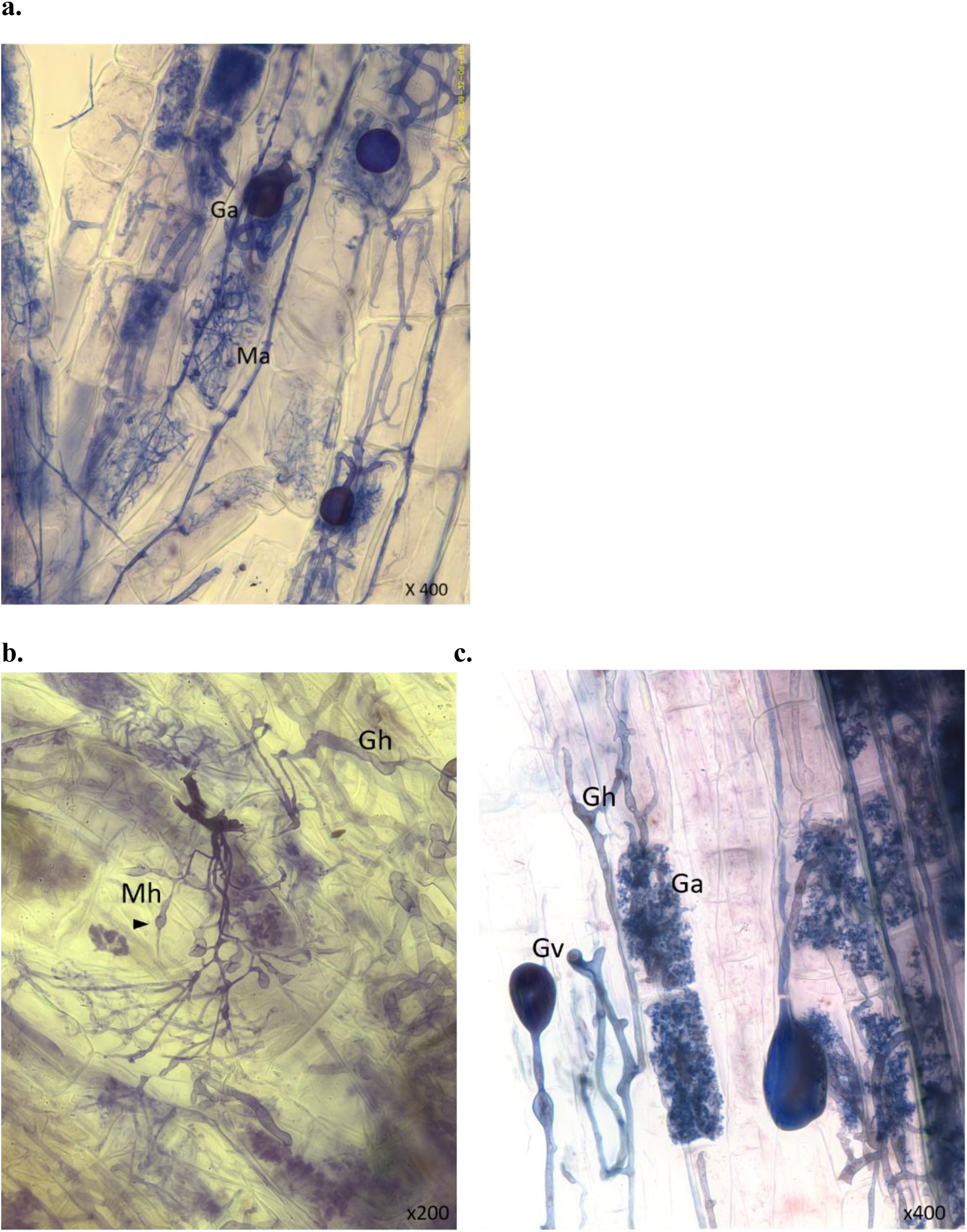
Microscopic observation of M-FRE and G-AMF. **a.** Distinct morphologies for M- (Ma) and G-arbuscules (Ga); **b.** Specific morphological features of M-FRE with swollen hyphae (Mh) and arbuscules surrounded by thicker G-AMF hyphae (Gh); **c.** Classical Glomeromycotina structures, with hyphae (Gh) and vesicles (Gv) next to arbuscules (Ga).

**Table 1.** Results of the linear mixed-effects ANOVA testing the effects of treatment (T), genotype (G), and their interaction (T × G) on root colonization (M%) by G-AMF and M-FRE.

| M % | Variable | Chisq | Df | Pr(>Chisq) |
| --- | --- | --- | --- | --- |
| G-AMF | Treatment | 0.178 | 1 | 0.673 |
|  | Genotype | 24.378 | 14 | **0.041 \*** |
|  | T x G | 13.535 | 14 | 0.485 |
| M-FRE | Treatment | 10.01 | 1 | **0.001\*\*** |
|  | Genotype | 12.431 | 14 | 0.571 |
|  | T x G | 10.500 | 14 | 0.72 |

**Figure 2.**
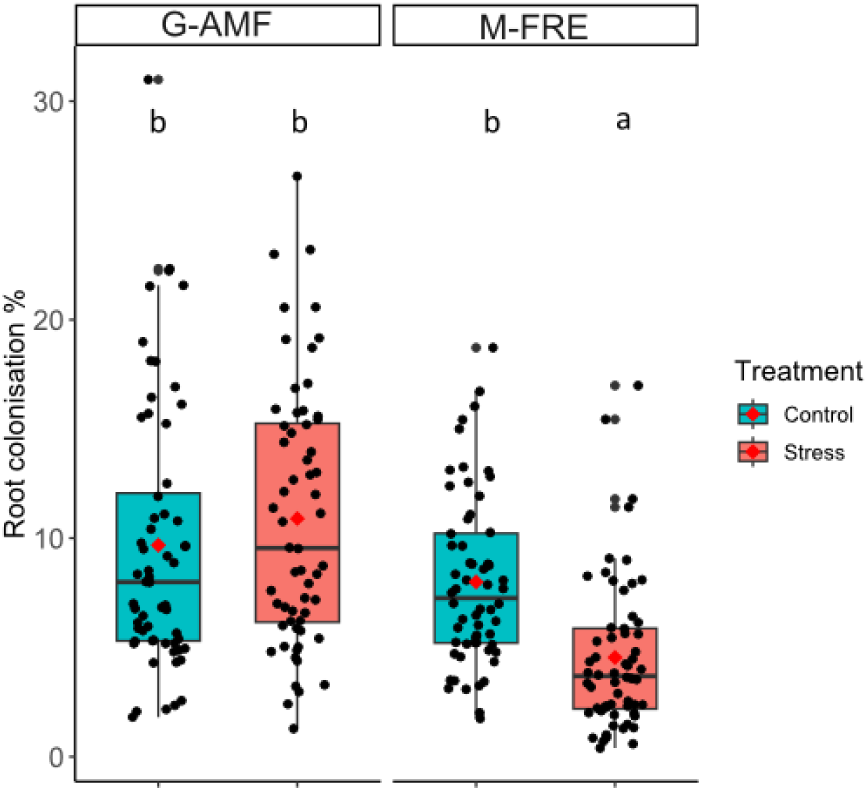
Colonization rate of durum wheat roots (all fifteen genotypes pooled) by G-AMF and M-FRE structures in the field experiment, under combined stress (reduced water and nitrogen) and control conditions. Boxplots represent the median and interquartile range, and red points indicate the mean. Letters above the boxplots indicate significant differences among colonization rates, as assessed by a post hoc Tukey test.

Durum wheat genotype had a significant impact on G-AMF colonization, but not on M-FRE. Concerning G-AMF, three varieties had significantly lower G-AMF colonization rates (Anvergur, RGT-Voilur, Casanova, Qualidou and Furio-Camillo with a mean colonization of about 7.3%, **Table S3 B**) than most genotypes. In contrast, the Kofa variety had the highest colonization rate (18.3 %).

Our approach, based on selecting the best-supported models from all possible combinations of predictors in generalized linear models (GLMs), allowed us to estimate standardized model coefficients and the relative importance of predictors in explaining variation in mycorrhization. Root traits and rhizosphere nutrient availability (N and P) emerged as key predictors of colonization, with contrasting associations for Glomeromycotina and Mucoromycotina (Fig. 3). The mean R² of the ten best-supported models was 0.207 for Glomeromycotina and 0.214 for Mucoromycotina. Colonization by the two mycorrhizal groups was positively associated, with relative importance values exceeding 1 for Mucoromycotina and 0.75 for Glomeromycotina.

**Figure 3.**
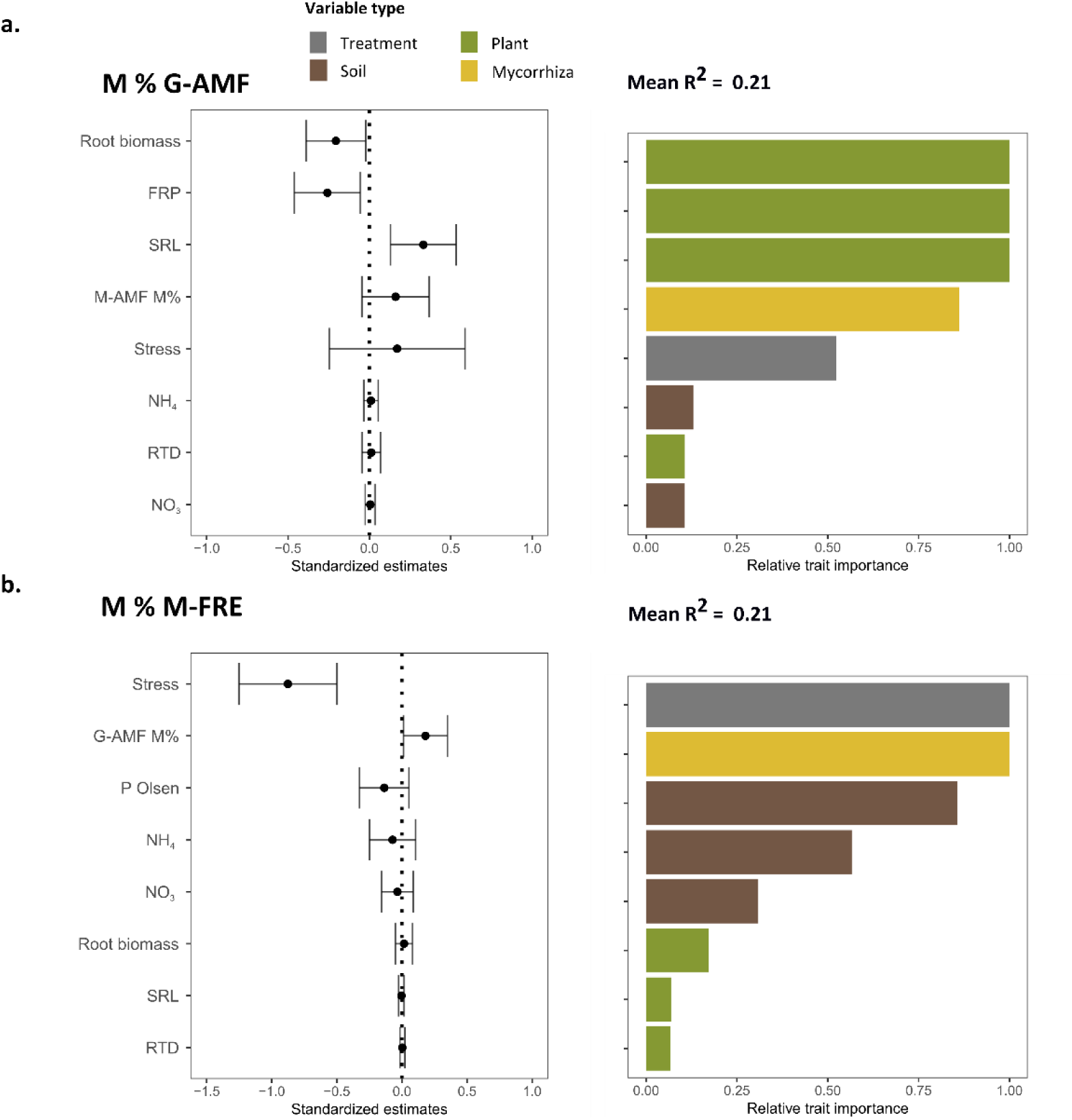
Standardized effects of traits on root colonization rates (M%) of Glomeromycotina (G-AMF, **a.**) and Mucoromycotina (M-FRE, **b.**) on fifteen durum wheat genotypes. For each mycorrhiza type, backward model selection was performed on a full model with colonization rate as the response variable and all traits as explanatory variables. Based on AICC, the top 10 models were retained to compute model-averaged estimates with their 95% unconditional confidence intervals. Plant traits are represented in green (FRP: fine root proportion, SRL: specific root length; RTD: root tissue density; root biomass), soil characteristics (nutrient availability in the rhizosphere: Olsen P, NH_4_, NO_3_) in brown and mycorrhizal colonization (of the other mycorrhizal type) in yellow and the treatment (control or combined water and nitrogen stress) in grey. Adjusted R-squared averaged across the top 10 models (R^2^) are reported for each modality.

For G-AMF, root traits consistently explained most of the variation in colonization. Colonization decreased with increasing root biomass and fine root production (FRP) and increased with specific root length (SRL) (**Fig. 3a**). Among soil variables, rhizosphere NO₃⁻ and NH₄⁺ were retained in the best-supported models (relative importance < 0.25; Fig. 3a), although the direction of their effects was inconsistent across models. The stress treatment was also included among the top models, but its effect was variable.

In contrast, Mucoromycotina colonization was primarily driven treatment effects and by soil variables. Stress exerted a consistently strong negative effect on colonization. Olsen P tended to show a negative association with Mucoromycotina colonization, while rhizosphere nitrate and ammonium exhibited weaker effects (**Fig. 3b**).

### Taxonomic composition and compartmental distribution of G-AMF and M-FRE

Overall, all OTUs belonged to nine families (**Fig. 4A**). Based on the SILVA classification, we identified a total of 74 OTUs assigned to the Glomeromycotina and 11 OTUs assigned to the Endogonaceae. Using the EUKARYOME taxonomic framework, nine of the Endogonaceae OTUs were assigned to the family recently renamed Planticonsortiaceae by Tedersoo *et al*. (2024). However, EUKARYOME failed to classify three additional Endogonaceae Taxa, which are likely M-FRE based on their phylogenetic position (**Fig. 5**) and BLAST similarity to sequences previously identified as putative MFRE. For this reason, we retained the SILVA taxonomic nomenclature throughout the manuscript. In addition, our internal BLAST against a curated reference database revealed one additional M-FRE sequence (Taxa 1337), resulting in a total of 12 Endogonaceae OTUs. The sequences of the 12 OTUs (Taxa 234, 27, 29, 1038, 1337, 1566, 429, 57, 15, 316) clustered with liverwort-associated Endogonaceae sequences, including MH174558, MH174557 and MH174584 (Rimington *et al.,* 2019; **Fig. 5**). Taxa 535 and 1928 grouped most closely with KX43777 (Orchard *et al.,* 2017a), a sequence shown to be identical to *Planticonsortium tenue* (Walker *et al.,* 2018). We therefore considered all 12 Endogonaceae Taxa identified in our dataset to represent M-FRE.

**Figure 4.**
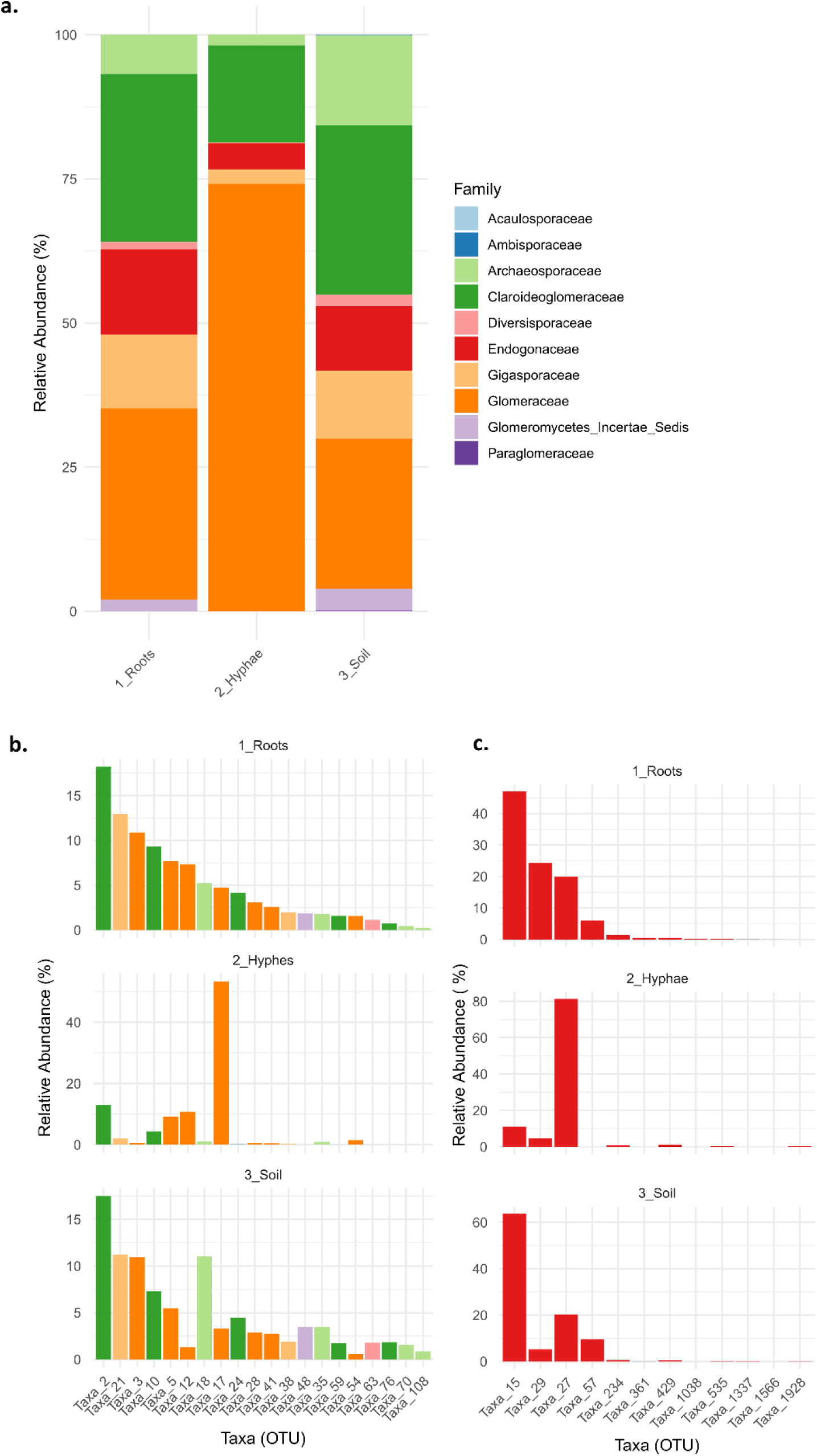
Relative abundance of mycorrhizal communities among the different compartments, in durum wheat roots DNA (Roots), extra-radical hyphae (Hyphae) and rhizospheric soil (Soil), at the Family level (**a.**), and focusing on the 20 most abundant OTUs of G-AMF (**b.**) and on all M-FRE OTUs (**c.**).

**Figure 5.**
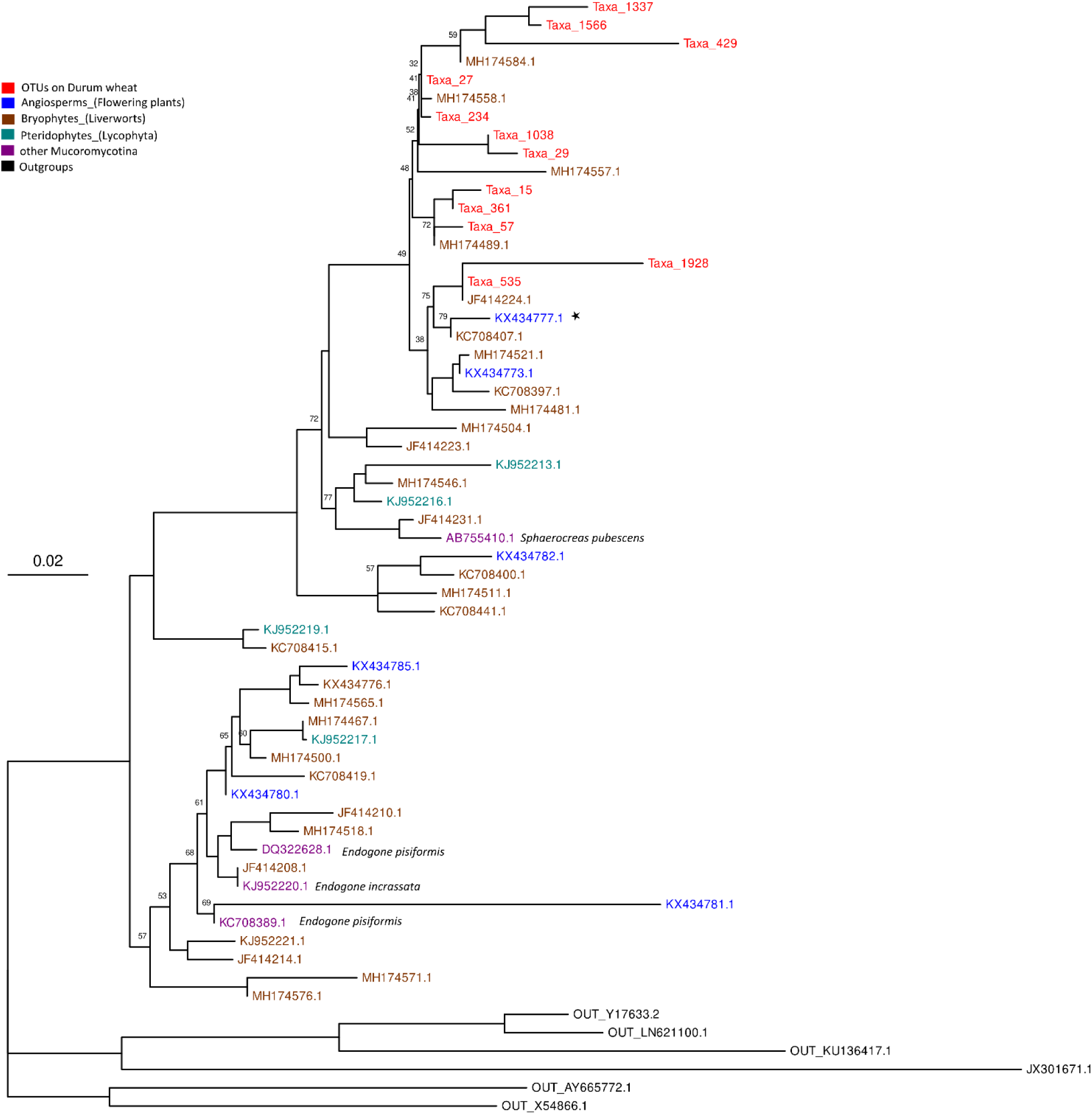
Phylogenetic tree of twelve Mucoromycotina OTUs sequenced from durum wheat roots (in red), built with IQ-TREE3. The tree includes fungal reference sequences, with the host plant indicated for each reference. Branch support values (ultrafast bootstrap approximation) are shown for values inferior to 80. The star marks the sequence corresponding to the reference isolate *Planticonsortium tenue*.

Among M-FRE sequences, OTU 15 was dominant in terms of relative abundance in root and soil samples, while OTU 27 was dominant in hyphal samples (**Fig. 4C**). The dominant taxa were largely shared among the three compartments, although their relative abundances differed. All OTUs were detected in roots, all except OTUs 1566 and 1038 were present in the soil, and five OTUs were not detected in the hyphal compartment. Within Glomeromycotina, Glomeraceae was the most abundant family both in terms of OTUs richness (31) and relative sequence abundance across root, hyphae, and soil samples (**Fig. 4A**). The 20 most abundant G-AMF OTUs accounted for 97% of all G-AMF sequences, and 97.5%, 98.3%, and 95.8% of sequences in roots, hyphae, and soil, respectively (**Fig. 4**). The most abundant OTU overall was OTU 2 (representing 15.4% of all sequences), assigned to Claroideoglomeraceae, which dominated in both root and soil samples, while OTU 17, a Glomeraceae displayed the highest relative abundance among the hyphae compartment.

No significant correlation was detected between root colonization by M-FRE and the number of M-FRE sequences (p = 0.313; R= 0.093). For G-AMF, a positive but weak correlation was observed between sequence number and colonization rate (p = < 0.001; R = 0.32).

### Stress effects on G-AMF and M-FRE diversity

As we wanted to test the global effect of stress on mycorrhizal communities in wheat, we focused on the combined community of roots and soil as hyphae sequencing depth was much lower, and weakening the data when rarefied for lower sample size. Regarding alpha-diversity, stress did not affect species richness (Hill 0) for either G-AMF or M-FRE (**Table 2 a.**), indicating that taxa were largely maintained. However, indices accounting for evenness and dominance (Hill 1 and Hill 2) were significantly reduced under stress for both fungal groups, reflecting shifts in relative abundances and a more uneven distribution of taxa.

**Table 2.**
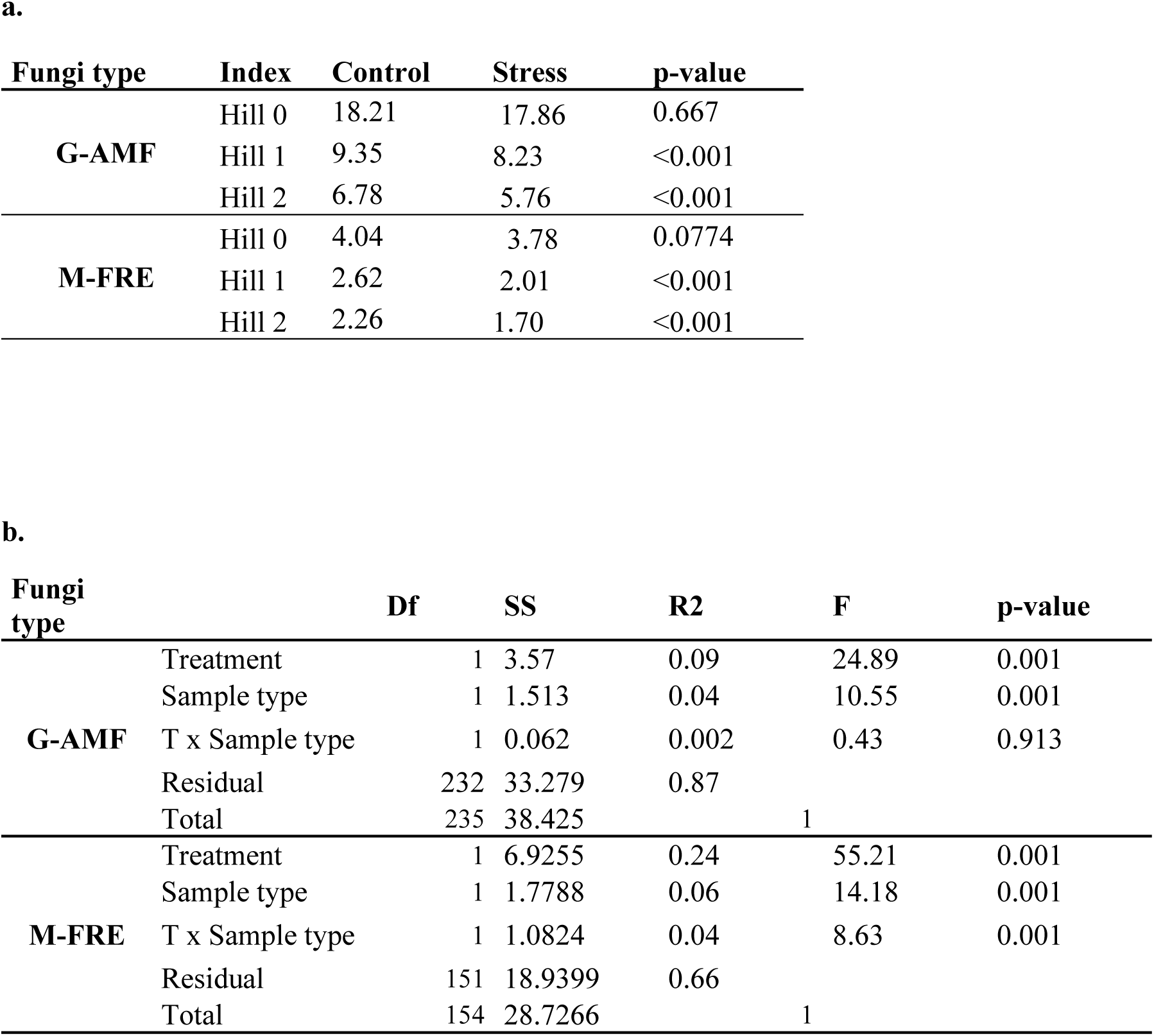
Diversity indices and PERMANOVA results for G-AMF and M-FRE communities. **a.** Mean Hill numbers (Hill 0 = species richness, Hill 1 = exponential of Shannon entropy, Hill 2 = inverse of Simpson concentration) for Glomeromycotina (G-AMF) and Mucoromycotina (M-FRE) under control and stress treatments. Differences between treatments were tested using Welch’s t-test. **b.** PERMANOVA testing the effects of stress treatment (T), sample type, and their interaction on community composition of G-AMF and M-FRE. Shown are degrees of freedom (Df), sum of squares (SS), proportion of variance explained (R²), F-statistic, and p-values.

For beta-diversity, both sample type and stress treatment affect mycorrhizal communities (**Table 2 b.**). In G-AMF communities, treatment was the main driver of community differences, explaining 24% of the variation, with additional effects of sample type (6%) and a significant treatment × sample type interaction (4%), indicating that treatment effects depended on sample type (**Table 2 b.**). In contrast, M-FRE communities showed weaker treatment (9%) and sample type (4%) effects, and no detectable significant interaction. Although PERMANOVA indicated higher treatment effects for M-FRE (**Table 2 b.**), caution is warranted: the M-FRE dataset contained only 12 taxa versus 74 for G-AMF, and lower taxonomic richness can inflate R² values. Betadisper tests showed no significant differences in multivariate dispersion for G-AMF (*p* = 0.26) and only a marginal, non-significant difference for M-FRE (*p* = 0.07), suggesting that the PERMANOVA results reflect genuine community differences rather than group spread.

## Discussion

Our study reveals contrasting responses of Mucoromycotina fine root endophytes (M-FRE) and Glomeromycotina arbuscular mycorrhizal fungi (G-AMF) to biotic and abiotic factors in durum wheat. Both fungal groups were consistently detected in roots, extraradical hyphae, and rhizospheric soil, often co-occurring within neighboring intraradical cells, suggesting that M-FRE not only colonize roots but also actively explore soil volumes. We further explored the phylogenetic diversity and host specificity of M-FRE, and how both fungal groups partitioned across compartments, potentially reflecting distinct functional strategies. Root colonization responded asymmetrically to stress and genotype: M-FRE colonization was strongly reduced under combined water and nitrogen limitation but remained insensitive to genotype, whereas G-AMF colonization was largely unaffected by stress yet varied significantly among wheat genotypes. Consistent with this, specific root length (SRL) and fine root proportion (FRP) were strongly correlated with G-AMF colonization, whereas no clear relationships emerged for M-FRE, highlighting fundamental differences in host–fungus coupling and raising questions of causality. Whether M-FRE relies on common symbiotic pathways remains uncertain, with contrasting evidence reported for Prout et al. (2024) and Cole et al. (2025). Similar ambiguity exists regarding autotrophic capabilities, pointing to a potentially wide continuum of ecological strategies within M-FRE.

Notably, under stress, M-FRE colonization was positively correlated with plant N and P uptake, whereas G-AMF colonization showed no such relationship, suggesting that M-FRE may play a key role in supporting plant nutrient acquisition. Finally, we examined how community diversity and the occurrence of specific taxa were reshaped by abiotic stress. These patterns frame the discussion of the ecological drivers and functional implications of M-FRE and G-AMF associations under field conditions, as further discussed below.

### Phylogeny and host range of M-FRE

We used the Sato *et al*. (2005) primers because, at the time of analysis, the Seeliger primers had not yet been published, although their use might have revealed greater M-FRE diversity (Seeliger *et al.,* 2024). The Endogonaceae sequences we detected were all clustered within clades that, in previous studies, have been consistently associated with the fine root endophyte (M-FRE) morphology. This allowed us, with caution, to refer to these sequences as M-FRE. It is noteworthy that one of our OTUs matched sequences originating from the M-FRE pot culture of the IBG reference strain BEG249 (International Bank for Glomeromycota, http://www.i-beg.eu/; matching KX434777.1 from Orchard *et al.,* 2017a), which was used for the formal description of the genus *Planticonsortium* (Walker *et al.,* 2018). Similarly, field studies conducted in Australia (Albornoz *et al.,* 2021) and in the United Kingdom (Seeliger *et al.,* 2024) reported that most of their mucoromycotinian OTUs clustered within the same clade containing sequences KX434773.1 and KX434777.1, which were originally dominant in strongly M-FRE-colonized roots of *Trifolium subterraneum* (Orchard *et al.,* 2017a). Our findings thus add a new record from southern France, indicating that taxa closely related to *Planticonsortium tenue* appear to be widespread globally. A better understanding of the diversity within *Planticonsortium* and the broader Densosporales (Lutz *et al.,* 2025) would be valuable, and future long-read sequencing could clarify which Mucoromycotina clades form M-FRE with arbuscules and their evolutionary relationships. Our M-FRE OTUs showed strong phylogenetic proximity to widespread OTUs found across diverse plant lineages, including non-vascular plants. This broad host range may support their role as host-generalist symbionts, suggesting evolutionary continuity with land plant–associated fungi, and is consistent with their widespread occurrence across most terrestrial biomes, as reported by Lutz *et al*. (2025).

### Niche partitioning and functional strategies across compartments

M-FRE OTUs were detected in all compartments—roots, rhizospheric soil, and hyphal bags—indicating active intra-radical colonization and potential extension into the surrounding soil. To our knowledge, this is the first metabarcoding-based evidence that most M-FRE, like G-AMF, occupy both intra- and extra-radical spaces. OTU 15 dominated roots and the rhizospheric soil, whereas OTU 27 was more frequent in hyphal bags, suggesting compartment-specific niche differentiation. The least abundant M-FRE taxa were absent from hyphal bags and soil, although low DNA yields from hyphal bags and the absence of quantitative abundance measures warrant cautious interpretation. It is also possible that the hyphal bag environment lacked cues or resources to attract all M-FRE or G-AMF taxa. The G-AMF communities showed strong compartment-specific patterns. Twenty dominant OTUs accounted for ∼97% of sequences, with OTU 2 being most abundant in roots and rhizospheric soil. In the hyphal compartment, OTU 17 represented more than 50% of the sequences, raising questions about its ecological strategy which possibly includes rapid soil exploration or sporulation under nutrient limitation. These patterns likely reflect differences in species life-history strategies—allocation to exploration versus host, sporulation biology, and stress responses—as previously discussed for G-AMF (Chagnon *et al.,* 2013). Functional traits and life-history perspectives are critical for understanding these dynamics, yet fine-scale ecological knowledge remains limited due to a lack of deployable trait-based tools beyond spore analysis. Hyphal bags provide a promising approach for broader-scale studies of intra-versus extra-radical investment, enabling direct observation of active mycelial and spores growth akin to root ingrowth bags. While Orchard *et al*. (2017a) reported a correlation between M-FRE colonization and sequence counts, and Albornoz *et al*. (2021) used M-FRE sequence abundance in roots as a proxy for colonization, our results indicate that this relationship is not consistent. Our findings are in line with those of Seeliger *et al*. (2024), despite their use of primers designed to encompass a broader diversity of M-FRE. For G-AMF, the weak correlation indicates that sequence counts may provide only a coarse estimate of fungal abundance. Overall, these findings underscore that sequence abundance should be interpreted with caution, and that root colonization is best assessed using microscopic observation.

### Abiotic drivers of root colonization by G-AMF and M-FRE under combined N and water stress

Overall root colonization of durum wheat was low (mean 16.8%) and appeared largely unaffected by combined water and N stress. When examined separately, G-AMF and M-FRE responded differently to abiotic and biotic factors. The M-FRE were particularly sensitive to limited water and N, which affected both colonization and diversity. The lack of host effects (genetic or traits) raises questions about plant control and whether M-FRE use the same molecular pathways for plant signaling and the same symbiotic pathways (Cole *et al.,* 2025; Prout *et al.,* 2024). While P availability did not affect G-AMF, even modest variation (13 ± 2 mg P Olsen) reduced M-FRE colonization, consistent with broader evidence that M-FRE are more P-sensitive than G-AMF. The tipping point for this response likely depends on plant genotype and critical P requirement (Jeffery *et al.,* 2018). In our field experiment, the strong reduction in M-FRE colonization under stress suggests that water limitation also played a key role. These patterns highlight the high sensitivity of M-FRE to interacting environmental gradients and the contrasting ecological strategies of these co-occurring root symbionts.

### Genotype-driven variability in G-AMF associations

Our results showed that, unlike M-FRE, G-AMF colonization was primarily shaped by plant genotype and root traits, including fine root proportion (FRP), specific root length (SRL), root tissue density, and root biomass. The positive correlation between high SRL and mycorrhization contrasts with the root economic spectrum, in which high SRL is linked to a “do-it-yourself” strategy (Bergmann *et al.,* 2020). While this concept has been developed across species, it was not confirmed here within the species of durum wheat. Some species, such as maize genotypes, appear to follow the root economic spectrum (Wild *et al.,* 2024). Relationships between mycorrhizal fungi and root traits remain poorly understood, and causality is unclear, as both plant and fungus can influence each other. Field measurements cannot disentangle causes and consequences. Few studies have investigated G-AMF effects on root morphology, and outcomes are species and genotype-dependent (de Souza Campos *et al.,* 2021). For example, colonization led to coarser roots in strawberries (Fan *et al.,* 2011) but increased fine root length in winter wheat (Lazarević *et al.,* 2018). In a pot experiment comparing M-FRE–dominated and mixed communities, average root diameter was up to 15% smaller in M-FRE-dominant treatments (Jeffery *et al.,* 2018).

### Context-dependent role of M-FRE in plant nutrient uptake

Root colonization did not necessarily correlate with plant nutrient uptake, and this was true for both N and P under most conditions. The only exception occurred under stress, where M-FRE but not G-AMF colonization correlated positively with nutrient content. However, this does not necessarily mean that M-FRE directly enhanced nutrient uptake; they might simply persist in plants that cope better with stress and acquire more N and P. Interestingly, M-FRE abundance and diversity declined under stress, yet they still correlated with nutrient content, suggesting that either efficient functioning or the activity of stress-tolerant taxa may be involved. To clarify the functional contribution of M-FRE, new isolates for controlled experimental testing will be necessary.

## Conclusion

In this study, G-AMF were primarily driven by biotic factors, whereas M-FRE were more responsive, and even sensitive, to abiotic conditions, particularly combined reduction in N and water availability. Previous work (Jeffery *et al.,* 2018) suggests that M-FRE colonization and function may vary strongly across plant species, potentially influencing their contributions to nutrient acquisition. Future studies should explore the complementarity of these two root symbioses, both in terms of plant-mediated regulation of colonization and their respective contributions to mycorrhizal functions, including nutrient uptake, immune system stimulation, the establishment of mycelial networks potentially capable of information transfer, and ultimately, their role in crop and soil health.

## Supporting information

Supplemental Results

## Author contribution

**Taschen E**.: Conceptualization, Formal analysis, Investigation, Methodology, Supervision, Visualization, Writing – original draft; **Guillot E.** : Conceptualization, Formal analysis, Investigation (Sampling, microscopy, soil analyses), Methodology, Supervision, Visualization, Data Curation, Writing – review and editing ; **Plassard C**.: Conceptualization, Writing – review and editing; **Kerbiriou E**.: Formal analysis, Writing – review and editing; **Dezette D**.: Investigation (sampling, molecular biology), Writing – review and editing; **Taudière A**.: Formal analysis, Writing – review and editing; **Personne A**.: Investigation (chemical analyses), **Robin A**.: Conceptualization, Investigation (molecular biology) Writing – review and editing; **Redecker D**.: Conceptualization, Writing – review and editing; **Hinsinger P**.: Funding acquisition, Conceptualization, Writing – review and editing.

## Acknowledgments

We thank Christine Tollon-Cordet (UMR AGAP) and the ARCAD laboratory (Montpellier) for their support with root DNA extractions, as well as Hélène Févrille and Pierre Roumet for their expertise on durum wheat and for setting up the field experiment. We also acknowledge the dedication and perseverance of the interns Marine Scellier and Arthur Houdaer. We thank the GEVES for field management, and ADNid—particularly Fabienne Moreau and Pascal Alonzo—for their efficient and constructive collaboration in carrying out the Illumina metabarcoding analyses. We thank Joshua Cole for comments on English and science.

## Sequencing data

Raw sequence data have been deposited in the European Nucleotide Archive (ENA) under accession number PRJEB107173.

