## Supplemental Results for "Contrasting biotic and abiotic drivers of Glomeromycotina and Mucoromycotina mycorrhizal associations in a durum wheat field"

**<u>Content :</u>**

**Supplementary Figure 1:** Field and sampling design

**Table S1:** Soil characteristics

**Table S2:** Durum wheat genotypes

**Table S3:** Mean colonization rates per genotype

**Supplemental Figure S2 :** Correlations plant nutrient uptake and colonisation rates.

**Supplementary Methods S1:** Root DNA extraction protocol

### Supplementary Figure 1

**a.**

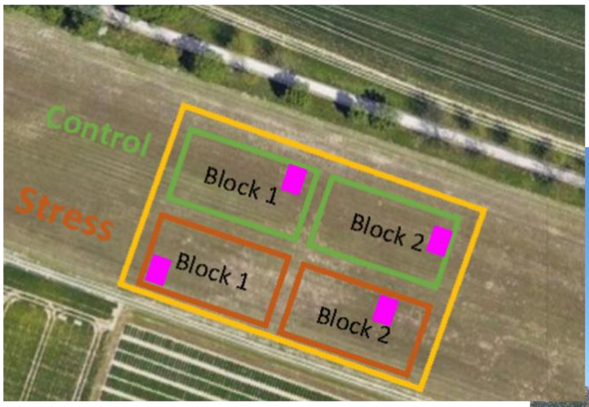


**b.**

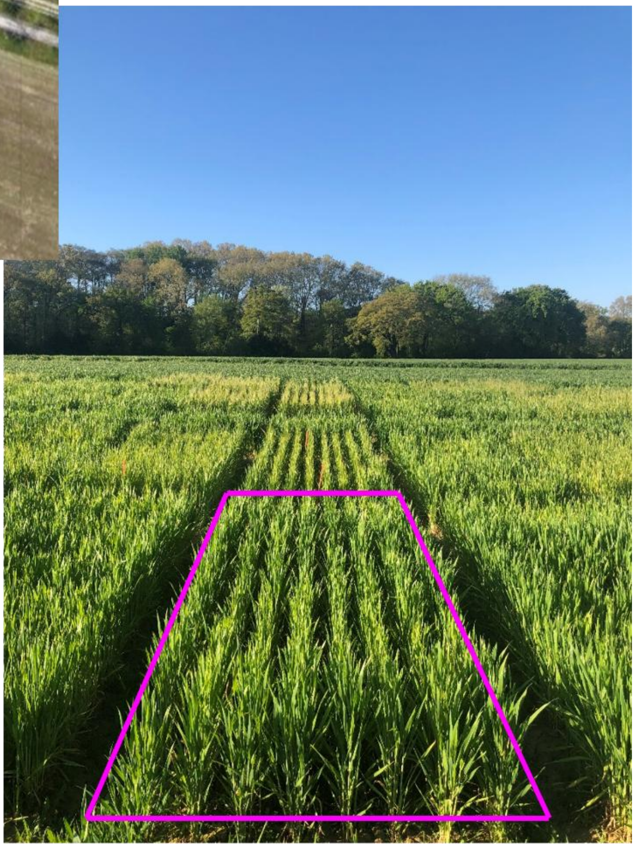

**a.** The field was divided in half (0.28 ha each) to apply two cultivation treatments): a **control** with optimal nutrient and water supply (151 units of N, two irrigations of 20 mm) and a combined water and nitrogen **stress** treatment (110 units of N, a single irrigation of 25 mm).

**b.** Wheat genotypes were cultivated on plots of 1.5 x 2.5 meters (**b.**), separated by a buffer of a common variety (RGT Voilur). Each genotype was repeated on two plots per treatments

**c.** One sampling unit corresponded to a single collection from a specific wheat genotype (ex: Genotype 1, in pink) within a plot. For each genotype and treatment, four samples were collected, from two plots (one per spatial block, a.), with each plot (b.) sampled at both the upper and lower positions (c.).

**c.**

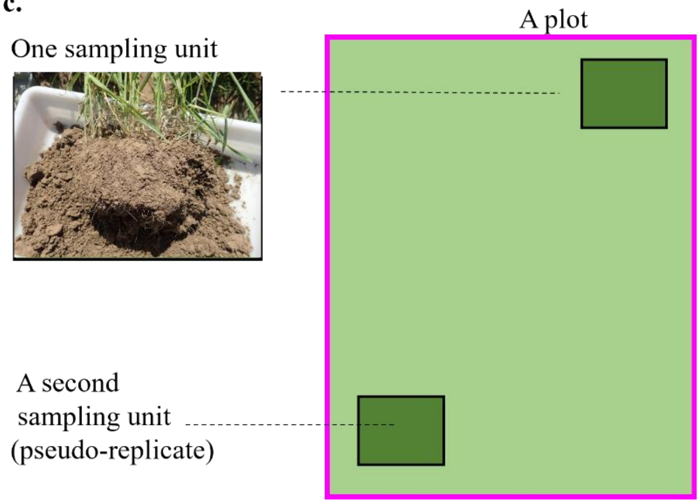

**d.** On each sampling unit we sampled and analyzed the following elements.

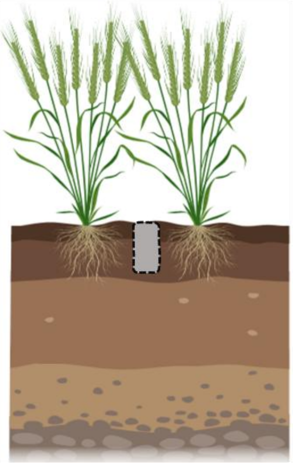

Shoot biomass, N and P content

Root biomass and **roots traits**

**DNA on 3 compartments** :
- roots
- rhizospheric soil
- hyphal compartment

Rhizospheric soil N – P availability

**e.** Photographs illustrating the installation of hyphal ingrowth bags. The nylon mesh had a pore size of 37 µm, and the sewn bags were filled with previously washed and dried coarse sand.

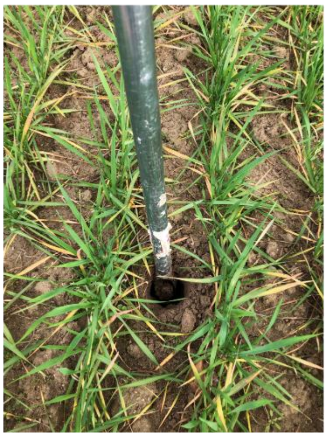

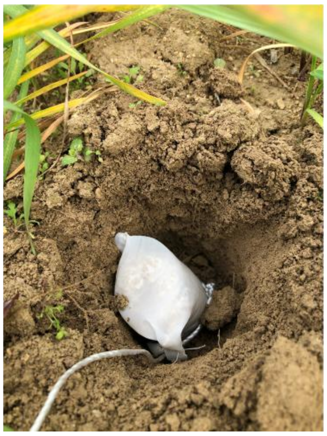

**Table S1**

**Soil characteristics** measured on 16 soil samples covering the experimental field.

| | unit | Mean | Minimum | Maximum |
|---|---|---|---|---|
| Clay (0 to 0,002 mm) | $g.kg^{-1}$ | 187,063 | 141,000 | 228,000 |
| Fine silt (0,002 to 0,02 mm) | $g.kg^{-1}$ | 331,500 | 235,000 | 400,000 |
| Coarse silt (0,02 to 0,05 mm) | $g.kg^{-1}$ | 140,000 | 121,000 | 165,000 |
| Fine sand (0,05 to 0,2 mm) | $g.kg^{-1}$ | 138,125 | 107,000 | 198,000 |
| Coarse sand (0,2 to 2,0 mm) | $g.kg^{-1}$ | 203,313 | 125,000 | 378,000 |
| Total nitrogen (N) | $g.kg^{-1}$ | 0,928 | 0,674 | 1,176 |
| Total calcium carbonate ($CaCO_3$) | $g.kg^{-1}$ | 731,708 | 658,000 | 796,000 |
| Total carbon (C) | $g.kg^{-1}$ | 95,914 | 88,684 | 100,964 |
| Total organic carbon (TOC) | $g.kg^{-1}$ | 8,109 | 5,444 | 10,501 |
| Carbon-to-nitrogen ratio (C/N) | - | 8,764 | 7,600 | 10,400 |
| Organic matter | $g.kg^{-1}$ | 14,033 | 9,420 | 18,200 |
| pH | - | 8,618 | 8,480 | 8,700 |
| Olsen phosphorus (P) | $g.kg^{-1}$ | 0,013 | 0,010 | 0,015 |
| Effective cation exchange capacity (CEC) | $cmol^{+}.kg^{-1}$ | 10,546 | 7,648 | 12,957 |
| Aluminium (Al) | $cmol^{+}.kg^{-1}$ | 0,028 | <0,02 | 0,038 |
| Calcium (Ca) | $cmol^{+}.kg^{-1}$ | 11,040 | 9,036 | 12,910 |
| Iron (Fe) | $cmol^{+}.kg^{-1}$ | 0,005 | 0,005 | 0,006 |
| Magnesium (Mg) | $cmol^{+}.kg^{-1}$ | 0,414 | 0,301 | 0,555 |
| Manganese (Mn) | $cmol^{+}.kg^{-1}$ | <0,005 | <0,005 | <0,005 |
| Potassium (K) | $cmol^{+}.kg^{-1}$ | 0,477 | 0,370 | 0,630 |
| Ammonium nitrogen ($N–NH_4^+$) | $mg.kg^{-1}$ | 2,049 | 1,290 | 2,557 |
| Nitrate nitrogen ($N–NO_3^-$) | $mg.kg^{-1}$ | 29,418 | 14,435 | 51,051 |
| Mineral nitrogen (Nmin) | $mg.kg^{-1}$ | 31,467 | 16,953 | 52,658 |

**Table S2**

Durum wheat genotypes, with year and origin of the inscription in official catalogues. Two of the most cultivated varieties in France were included, with anvergur being the dominant variety representing 55.7% of the surfaces cultivated with durum wheat, and RGT VOILUR representing 9.4 % (ARVALIS, 2025).

| Durum wheat Genotypes | Inscription date in the catalogue | Inscription in original catalogues (Country or Institution) |
|---|---|---|
| ACADUR | 2015 | Italy |
| ASTERIX | 2012 | Italy |
| AZEGHAR-2_DP128 | 1992 | Icarda |
| KOFA | 1995 | USA |
| COLOSSEO_DP087 | 1995 | Italy |
| QUALIDOU | 2011 | France |
| FURIO-CAMILLO | 2012 | Italy |
| ANTALIS | 2013 | Italy |
| CASANOVA | 2003 | Italy |
| MONASTIR | 2012 | France |
| ANVERGUR | 2012 | France |
| NEMESIS | unknown | |
| GIUSTO | 2001 | Italy |
| EL4X_35 | non cultivated, EPO[1] | |
| RGT_VOILUR | 2016 | France |

[1] EPO: Evolutionary Pre-breeding population (David et al., 2014)

David J, Holtz Y, Ranwez V et al (2014) Genotyping by sequencing transcriptomes in an evolutionary pre-breeding durum wheat population. Mol Breed 34:1531–1548

**Table S3**

Mean colonization percentage (emmeans {emmeans}) of G-AMF (a.) and M-FRE (b.) on the different wheat genotypes including all (both control and stress treatment), shown here without arcsinus transformation for ease of interpretation.

**a.** G-AMF

| Genotype | Emmean | SE | df | lower.CL | upper.CL | .group |
|---|---|---|---|---|---|---|
| anvergur | 6.844750 | 1.932662 | 90 | 3.005179 | 10.68432 | a |
| casanova | 6.957868 | 1.932662 | 90 | 3.118297 | 10.79744 | a |
| rgt_voilur | 6.960063 | 1.932662 | 90 | 3.120491 | 10.79963 | a |
| qualidou | 7.330650 | 1.932662 | 90 | 3.491079 | 11.17022 | a |
| furio_camillo | 8.238938 | 1.932662 | 90 | 4.399366 | 12.07851 | a |
| azeghar | 9.244427 | 1.932662 | 90 | 5.404856 | 13.08400 | ab |
| acadur | 9.607775 | 1.932662 | 90 | 5.768204 | 13.44735 | ab |
| nemesis | 10.225692 | 1.932662 | 90 | 6.386121 | 14.06526 | ab |
| monastir | 11.235375 | 1.932662 | 90 | 7.395804 | 15.07495 | ab |
| el4x_35 | 11.298639 | 1.932662 | 90 | 7.459068 | 15.13821 | ab |
| asterix | 11.394200 | 1.932662 | 90 | 7.554629 | 15.23377 | ab |
| antalis | 11.669075 | 1.932662 | 90 | 7.829504 | 15.50865 | ab |
| colosseo | 12.398350 | 1.932662 | 90 | 8.558779 | 16.23792 | ab |
| giusto | 12.666175 | 1.932662 | 90 | 8.826604 | 16.50575 | ab |
| kofa | 18.275597 | 1.932662 | 90 | 14.436026 | 22.11517 | b |

**b.** M-FRE

| Genotype | emmean | SE | df | lower.CL | upper.CL | .group |
|---|---|---|---|---|---|---|
| rgt_voilur | 4.502437 | 1.332301 | 90 | 1.855589 | 7.149286 | a |
| nemesis | 4.561962 | 1.332301 | 90 | 1.915114 | 7.208811 | a |
| azeghar | 4.856125 | 1.332301 | 90 | 2.209276 | 7.502974 | a |
| casanova | 4.916675 | 1.332301 | 90 | 2.269826 | 7.563524 | a |
| qualidou | 5.131850 | 1.332301 | 90 | 2.485001 | 7.778699 | a |
| anvergur | 5.207125 | 1.332301 | 90 | 2.560276 | 7.853974 | a |
| kofa | 5.974750 | 1.332301 | 90 | 3.327901 | 8.621599 | a |
| acadur | 6.192225 | 1.332301 | 90 | 3.545376 | 8.839074 | a |
| colosseo | 6.346100 | 1.332301 | 90 | 3.699251 | 8.992949 | a |
| furio_camillo | 6.851312 | 1.332301 | 90 | 4.204464 | 9.498161 | a |
| el4x_35 | 7.326500 | 1.332301 | 90 | 4.679651 | 9.973349 | a |
| asterix | 7.533800 | 1.332301 | 90 | 4.886951 | 10.180649 | a |
| antalis | 8.070425 | 1.332301 | 90 | 5.423576 | 10.717274 | a |
| monastir | 8.126012 | 1.332301 | 90 | 5.479164 | 10.772861 | a |
| giusto | 8.446325 | 1.332301 | 90 | 5.799476 | 11.093174 | a |

### Supplemental Table S4

Mean values for variables measured on wheat plant shoots per treatment and comparisons between treatments (t-test and Bonferroni correction). Units are standardized.

| Variable | Unit | Mean (Stress) | Mean (NoStress) | P-value |
|---|---|---|---|---|
| Shoot biomass | g/plant | 1.230 | 1.621 | <0.001 |
| N concentration | % DW | 0.909 | 1.436 | <0.001 |
| P concentration | % DW | 0.171 | 0.187 | <0.001 |
| Total N per plant | mg/plant | 10.93 | 22.99 | <0.001 |
| Total P per plant | mg/plant | 2.071 | 2.995 | <0.001 |

### Supplemental Figure S2

Pearson correlations between mean plant P uptake (a., b.) and N uptake (bC., d.) and root colonization by G-AMF (a., c.) and M-FRE (b., d.) for each treatment (stress : red ; control : blue).

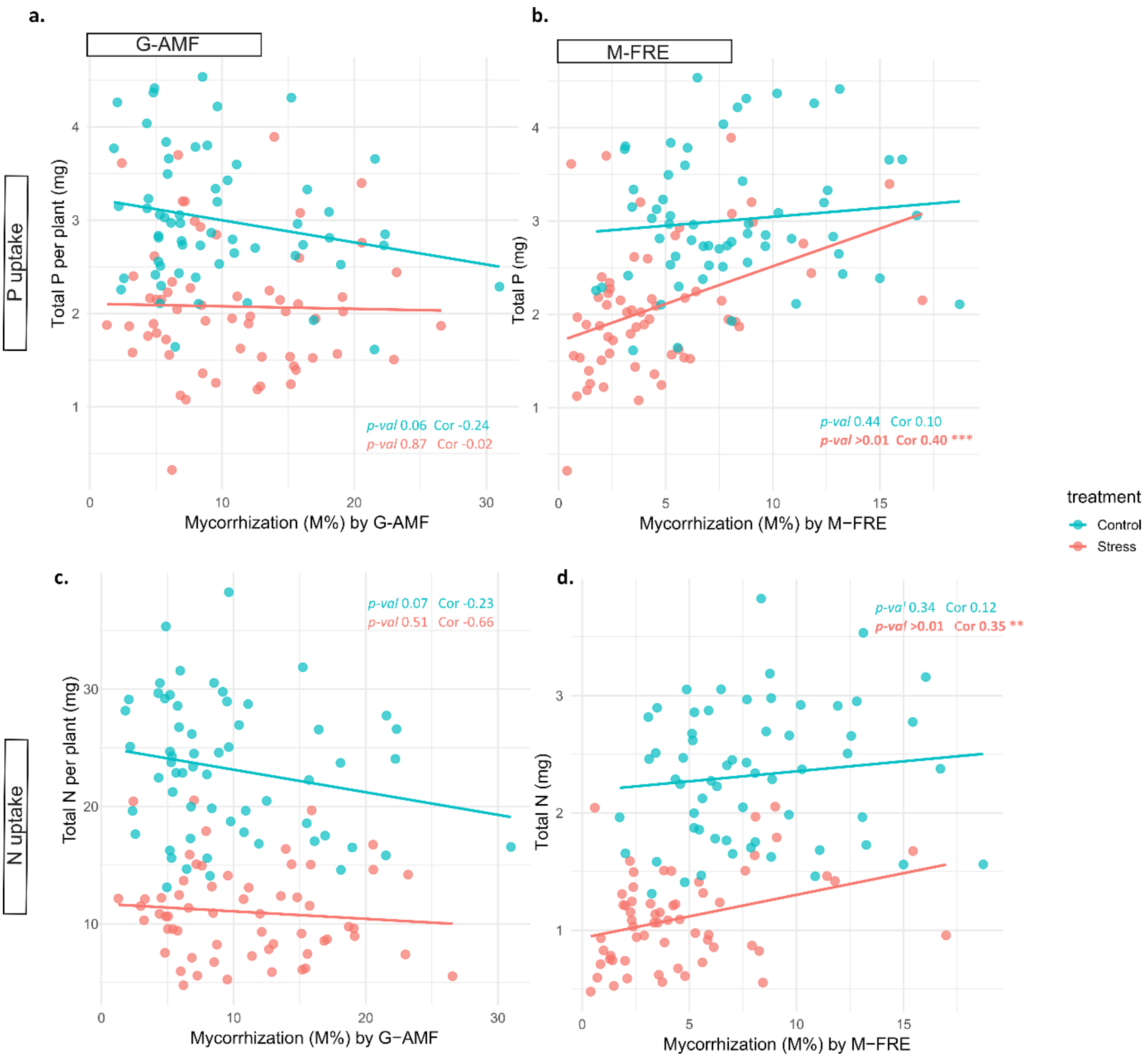

**Supplementary Methods S1**

**Root DNA extraction (MATAB lysis + magnetic-bead purification on KingFisher™ Flex)**
Developped by Christine Tollon (UMR ARCAD, INRAE,Montpellier) and Damien Dezette (UMR Eco&Sol, INRAE, Montpellier)

**Overview**

Genomic DNA was extracted from cryo-ground roots using a hot MATAB lysis with RNase treatment, followed by magnetic-bead binding and automated purification on a KingFisher™ Flex. Final elution was in TE 1×.

---

**Reagents and solutions**

**Lysis buffer (MATAB; prepare fresh or same day)**

For **50 mL**:

- Tris-HCl, **1 M**, pH 8.0 … **5 mL**
- NaCl, **5 M** … **14 mL**
- EDTA, **0.5 M**, pH 8.0 … **2 mL**
- MATAB (mixed alkyltrimethylammonium bromide) … **0.990 g**
- PEG-6000 … **0.495 g**
- $Na_2SO_3$ (sodium sulfite, antioxidant) … **0.253 g**
- Nuclease-free water … **to 50 mL total**

**Preparation:** Combine salts/buffers, dissolve PEG and MATAB with gentle warming. Add $Na_2SO_3$ last. Bring to volume, **preheat to 72 °C** before use.

Note: We used **500 µL** MATAB per sample (instead of 400 µL) to ensure complete pellet resuspension.

**Other reagents**

- **RNase A** (e.g., 10 mg/mL stock). Add **2 µL** per sample immediately after MATAB addition.
- **Guanidinium chloride, 7.8 M** (binding additive)
- **Isopropanol**, molecular biology grade
- **Magnetic beads** suitable for DNA binding (as per AGAP/KingFisher protocol)
- **Wash buffers 1–5** (see KingFisher program notes below; 600 µL per well each)
- **TE 1×** (10 mM Tris-HCl, 1 mM EDTA, pH 8.0), for elution

**Consumables**

- 2.0 mL screw-cap tubes (for grinding)
- 96-well deep-well plates (compatible with KingFisher; ≥1 mL per well) × **5** for washes
- 96-well deep-well plate for **binding**
- 96-well elution plate (standard height)
- 96-well storage plate
- KingFisher tip combs (as required by program)
- Cryovials for ground tissue
- **Liquid nitrogen**
- **70% ethanol** (for tool decontamination)

---

**Equipment**

- Cryogenic grinder (or equivalent) with: **1 large steel bead + 6 bicone beads** per tube
- Vortex mixer
- Temperature-controlled incubator/thermomixer set to **72 °C** with horizontal agitation
- Microcentrifuge capable of **12,500 rpm**, room temperature
- **KingFisher™ Flex** (Thermo Fisher Scientific)
- Magnetic stand for 96-well plates
- Plate centrifuge (capable of **2,000 rpm**)
- Analytical balance
- –80 °C freezer

---

**Sample preparation (cryo-grinding)**

1. **Pre-chill** tools and tubes with **liquid nitrogen**.

2. Aliquot ~**100 mg** root tissue into 2.0 mL tubes. Keep submerged/on $LN_2$.
3. Add **1 large bead + 6 bicone beads** to each tube.
4. **Grind** at **1100 rpm for 1 min**; **repeat 2–3×** as needed to obtain fine powder.
5. Keep ground material cold (liquid nitrogen) and store at **–80 °C** until extraction.
6. **Decontamination:** Clean spatulas/tools with **70% ethanol** between samples.

Throughput tip: Process **≤24 tubes** per batch to maintain timing and temperature control.

---

**DNA extraction (hot MATAB + RNase)**

For each batch (≤24 samples):

1. **Preheat** MATAB buffer to **72 °C**.
2. Retrieve tubes with **100 mg** frozen root powder from –80 °C.
3. Add **500 µL** hot MATAB to each tube.
4. Immediately add **2 µL RNase A** to each tube.
5. **Invert** each tube once to wet all powder; **vortex** briefly.
6. **Incubate 1 h at 72 °C**, tubes **horizontal**, under agitation.
7. **Centrifuge** at **12,500 rpm for 15 min** at room temperature.
8. Carefully transfer **350 µL supernatant** from each tube to the **Binding plate** (see next section).

---

**Magnetic-bead binding and KingFisher™ Flex purification**

**Prepare the Binding plate (96-well deep-well)**

Per well (prepare the number of wells matching your samples):

- **160 µL** guanidinium chloride **7.8 M**
- **450 µL** isopropanol
- **15 µL** magnetic beads
  Mix by gentle tapping or pipetting.

Add the **350 µL lysate supernatant** (from step above) directly into each prepared well. Seal or cover to prevent evaporation.

**Prepare Wash and Elution plates**

- **Wash 1–5 plates:** 96-well deep-well plates with **600 µL** of **Wash buffer 1**, **2**, **3**, **4**, and **5** respectively.
- **Elution plate:** 96-well plate with **100 µL TE 1×** per well.

**Program dependency:** Wash buffer compositions follow the AGAP/KingFisher method **PEX-EXT-002q** (a lab protocol that defines the five sequential washes). If using an equivalent program, match plate order and volumes accordingly.

**KingFisher run**

1. Load plates and tip comb(s) per the deck map required by **PEX-EXT-002q**.
2. Start program **PEX-EXT-002q**. The instrument will perform binding, wash steps 1–5, and transfer to elution.
3. After the run, **centrifuge the elution plate** at **2,000 rpm for 2 min** to sediment residual beads.
4. Place elution plate on a **magnetic stand** and transfer the supernatant (DNA) to a **storage plate**.
5. **Store at 4 °C** short-term (same day) or **–20 °C/–80 °C** for long-term.

---

**Quality control**

- **Concentration:** Quantify DNA by fluorometry (e.g., Qubit dsDNA HS or PicoGreen) or Nanodrop
- **Purity:** Record A260/280 and A260/230 ratios if using spectrophotometry.
- **Integrity:** Run **1% agarose gel** or capillary electrophoresis on a subset.

---

**Notes and troubleshooting**

- **Pellet resuspension:** Using **500 µL** MATAB (vs. 400 µL) improved recovery from certain root matrices.

- **RNase volume:** Add **immediately after** lysis buffer; delayed RNase can increase carryover RNA.
- **Carryover beads:** If beads remain after elution, repeat the **2,000 rpm, 2 min** spin and magnetic clearing before transfer.
- **Throughput:** Keep batches at **≤24** to maintain timing consistency.
- **Contamination control:** Clean tools with **70% ethanol** between samples; maintain a cold chain during grinding.
- **Program compatibility:** If **PEX-EXT-002q** is unavailable, use an equivalent KingFisher method with: (i) guanidinium/isopropanol binding, (ii) **five sequential washes** (composition matching your validated SOP), (iii) **TE 1×, 100 µL** elution.

---

**Safety**

- Handle **hot (72 °C)** buffers with heat-resistant gloves.
- Use appropriate PPE when handling **guanidinium salts**, **isopropanol**, and **liquid nitrogen**.
- Dispose of chemical and biological waste according to institutional guidelines.

> Indications:

On this project mean DNA yield per sample was 62 ng/µl (SE 26 ng/µ; max: 184; min : 17 ng/µl).
